# Context-dependent functions of mitochondria protein quality control in lung

**DOI:** 10.1101/2022.12.08.519642

**Authors:** Le Xu, Chunting Tan, Justinn Barr, Nicole Talaba, Jamie Verheyden, Ji Sun Chin, Samvel Gaboyan, Ira Advani, Ruth M Elgamal, Kyle J Gaulton, Grace Lin, Kamyar Afshar, Eugene Golts, Angela Meier, Laura E Crotty Alexander, Zea Borok, Yufeng Shen, Wendy K Chung, David J McCulley, Xin Sun

## Abstract

Aside from its role as the universal energy source of the cell, mitochondria also control many aspects of cell behavior. In an intact tissue, whether all cells require mitochondria function to the same extent, and how mitochondria insufficiency impacts cell behavior are poorly understood. Here we show that in the mouse lung epithelium, inactivation of LONP1, an energy ATP-dependent protease that functions in the mitochondria to degrade unfolded and misfolded proteins, led to mitochondria deficiency. In the naïve epithelium of the developing lung, loss of *Lonp1* obliterated cell proliferation and differentiation. In the adult airway epithelium during homeostasis, loss of *Lonp1* led to selective death of terminally differentiated multiciliated cells, leading to a cascade of progenitor activation to replace lost cells. In the adult airway epithelium following influenza infection, loss of *Lonp1* led to failure of airway progenitor migration into the damaged alveolar region. Bulk and single cell transcriptomic analysis revealed that one branch of the ER stress pathways, namely integrated stress response (ISR), is ectopically upregulated in mutants under all three conditions. Inactivation of core ISR transcription factor ATF4 in the *Lonp1* mutant airway reversed abovementioned phenotypes. Taken together, our findings demonstrate that depending on a cellular context, intact mitochondria function is required in either progenitor or progeny cells, and is essential for cell proliferation, survival or migration in the mammalian lung.

## INTRODUCTION

The human airway is composed of a colorful array of cell types each with assigned function (Cardoso and Lu, 2006; Morrisey and Hogan, 2010; Tata and Rajagopal, 2017) These include, among many other relatively rare cell types, luminal club cells that moisturize the air, luminal ciliated cells that keeps the airway clean, and basal progenitor cells that lay in reserve to replace lost luminal cells. A balanced ratio of these cell types is essential for proper airway function. Aside from its role in conducting air and serving as a barrier against airborne particles and pathogens, the airway epithelium is also a source of cells that can respond to severe alveolar injury (Kathiriya et al., 2020; Kumar et al., 2011; Vaughan et al., 2015). How the proper airway cell type balance is established, maintained and restored remains poorly understood.

As a part of the ancestral endomembrane systems, mitochondria are present in all eukaryotic cells. Aside from their canonical roles as the powerhouse of the cell that convert oxygen and nutrients into energy in the form of ATP (Kuhlbrandt, 2015), mitochondria dynamically modulate many other fundamental cellular processes, such as cell metabolism (Spinelli and Haigis, 2018), reactive oxygen species balance (Murphy, 2009) and calcium homeostasis (Duchen, 2000). Mitochondria also controls many aspects of cell behaviors, including cell survival, proliferation, differentiation and movement (Lisowski et al., 2018; Paupe and Prudent, 2018; Wang, 2001). As a result, mitochondria dysfunction is implicated in the pathogenesis of many human disorders, including lung diseases such as chronic obstructive pulmonary disease (COPD), asthma and idiopathic pulmonary fibrosis (Aghapour et al., 2020; Mabalirajan et al., 2008; Prakash et al., 2017). In lung, recent studies revealed that mitochondria-mediated calcium homeostasis in alveolar type II cells (AT2s), the facultative progenitors in the alveolar epithelium, play essential roles in the process of surfactant secretion and cell differentiation post injury (Ali et al., 2022; Islam et al., 2022). Decrease of mitochondrial abundance, as well as perturbations of their intracellular distribution within AT2s, have also been shown to impair key signaling during alveologenesis (Zhang et al., 2022). How mitochondria activity contribute to airway epithelial cell fate decisions under normal and injured condition is largely unknown. Given their dynamic cellular composition and constant exposure to environmental insults, the airway epithelium may serve as a rich setting to explore mitochondria regulated cell behaviors.

Precise synthesis, folding and assembly of thousands of proteins in the mitochondria is essential for their health and function. Mitochondria protein quality is closely monitored by a group of proteases that are specialized for degrading unfolded, misfolded or damaged protein substrates (Quiros et al., 2015). AAA+ Lon protease 1 (LONP1) is a serine peptidase that is evolutionarily conserved from bacteria to human. In eukaryotes, LONP1 homo-oligomerizes to form a soluble hexameric ring in the mitochondria matrix (Shin et al., 2021b; Yang et al., 2022), where it binds and cleaves a broad range of substrates, including, but not limited to, damaged components of the electron transport chain (ETC), mitochondrial enzymes aconitase, cytochrome c oxidase COX4-1, steroidogenic acute regulatory protein and 5-aminolevulinic acid synthase, mitochondrial transcription factors TFAM and MTERF2 and the DNA polymerase POLG (Bota and Davies, 2002; Fukuda et al., 2007; Ghosh et al., 2019; Granot et al., 2007; Lu et al., 2013; Matsushima et al., 2010; Silva-Pinheiro et al., 2021; Tian et al., 2011; Zurita Rendon and Shoubridge, 2018). In addition, LONP1 was found to possess protease-independent chaperone activity in protein stabilization (Shin et al., 2021a), as well as the DNA binding activity in mitochondrial DNA maintenance (Chen et al., 2008; Fu and Markovitz, 1998; Lu et al., 2003). Consistent with its multifaceted roles in maintaining mitochondria health, a decrease in *Lonp1* transcripts is associated with common diseases such as Parkinson’s disease (Sanchez-Lanzas and Castano, 2021). An array of *Lonp1* mutations are associated with rare disorders such as cerebral, ocular, dental, auricular, skeletal (CODAS) syndrome (Strauss et al., 2015) and congenital diaphragmatic hernia (CDH) (Qiao et al., 2021).

Global inactivation of *Lonp1* in mice causes early embryonic lethality during early gastrulation (Quiros et al., 2014). Inactivation of *Lonp1* in skeletal muscles led to autophagy and muscle loss (Xu et al., 2022), but protected the mice from diet-induced obesity through a long-range metabolic response (Guo et al., 2022); in oocytes led to premature ovarian insufficiency and infertility (Sheng et al., 2022); in cardiac progenitors led to defective ventricular development (Zhao et al., 2022); in adult cardiomyocytes led to dilated cardiomyopathy (Lu et al., 2019). These findings indicate a crucial role of LONP1 in organ development and homeostasis, which has not been studied in lung.

Here, we show that inactivation of *Lonp1* in the developing lung epithelium, but not the lung mesenchyme, led to a profound defect in the proliferation and differentiation of the naïve epithelial cells. In comparison, inactivation of *Lonp1* in the adult airway during homeostasis led to selective apoptosis in terminally differentiated cells, which led to mobilization of the club and basal progenitors to replace these lost cells. Following influenza injury, inactivation of *Lonp1* led to inability of the airway progenitors to migrate into the damaged alveoli. We found that the integrated stress response (ISR) pathway is ectopically activated in the *Lonp1* mutants, and contributes to the phenotypes. Our findings demonstrate that mitochondria proteostasis is rate limiting in different cell types, and plays an array of essential roles in lung epithelial cells throughout their lifespan.

## RESULTS

### LONP1 controls epithelium proliferation, signaling and branching morphogenesis in the developing lung

To first investigate the requirement for LONP1 during lung development, we assessed its expression in scRNA-seq datasets of the embryonic mouse lung (Zepp et al., 2021). *Lonp1* is broadly expressed at similar levels in all lung compartments, including epithelial, mesenchymal, endothelial and immune cells (Figure S1A), suggesting that LONP1 may play important roles in multiple cell lineages. To test this hypothesis, we inactivated *Lonp1* in selected lineages by generating either *Tbx4rtTA;tetOcre;Lonp1^flox/flox^* (hereafter *Tbx4rtTA;tetOcre;Lonp1*) or *Shh^cre/+^;Lonp1^flox/flox^* (hereafter *Shhcre;Lonp1*) mice. In *Tbx4rtTA;tetOcre;Lonp1* mice, continuous doxycycline administration starting at E6.5, know to trigger cre activity broadly in the lung mesenchyme and endothelium, led to an efficient reduction of *Lonp1* transcripts (∼70% reduction, Figure S1B). Mutant mice are viable with no obvious abnormalities in lung by histology (Figure S1C). These results indicate that LONP1 is dispensable in mesenchymal and endothelial cells for lung development.

In *Shhcre;Lonp1* mutants, the cre-mediated recombination led to efficient reduction of *Lonp1* transcripts ((∼50% reduction, Figure S1D). Mutants were lethal at birth. Tracing back to embryonic day (E) 13.5, the mutant lungs were exhibited reduced tip number and dilated tips (Figures 1A and 1B). Wholemount immunostaining of E-Cadherin to highlight the epithelium showed that these defects manifested as early as E11.5, at the start of secondary branching (Figure 1C). To address if *Lonp1* plays a role in lung proximal-distal patterning as well, we examined the expression patterns of *Sox2* (marker for proximal epithelium) and *Sox9* (marker for distal epithelium) at both the RNA and protein levels. There is a reduction of *Sox2* domain and expansion of *Sox9* domain in the mutant compared to control (Figures S1E-S1G). Taken together, these results suggest that LONP1 is essential in the lung epithelium for branching morphogenesis and proximal-distal epithelium patterning.

**Figure 1.**
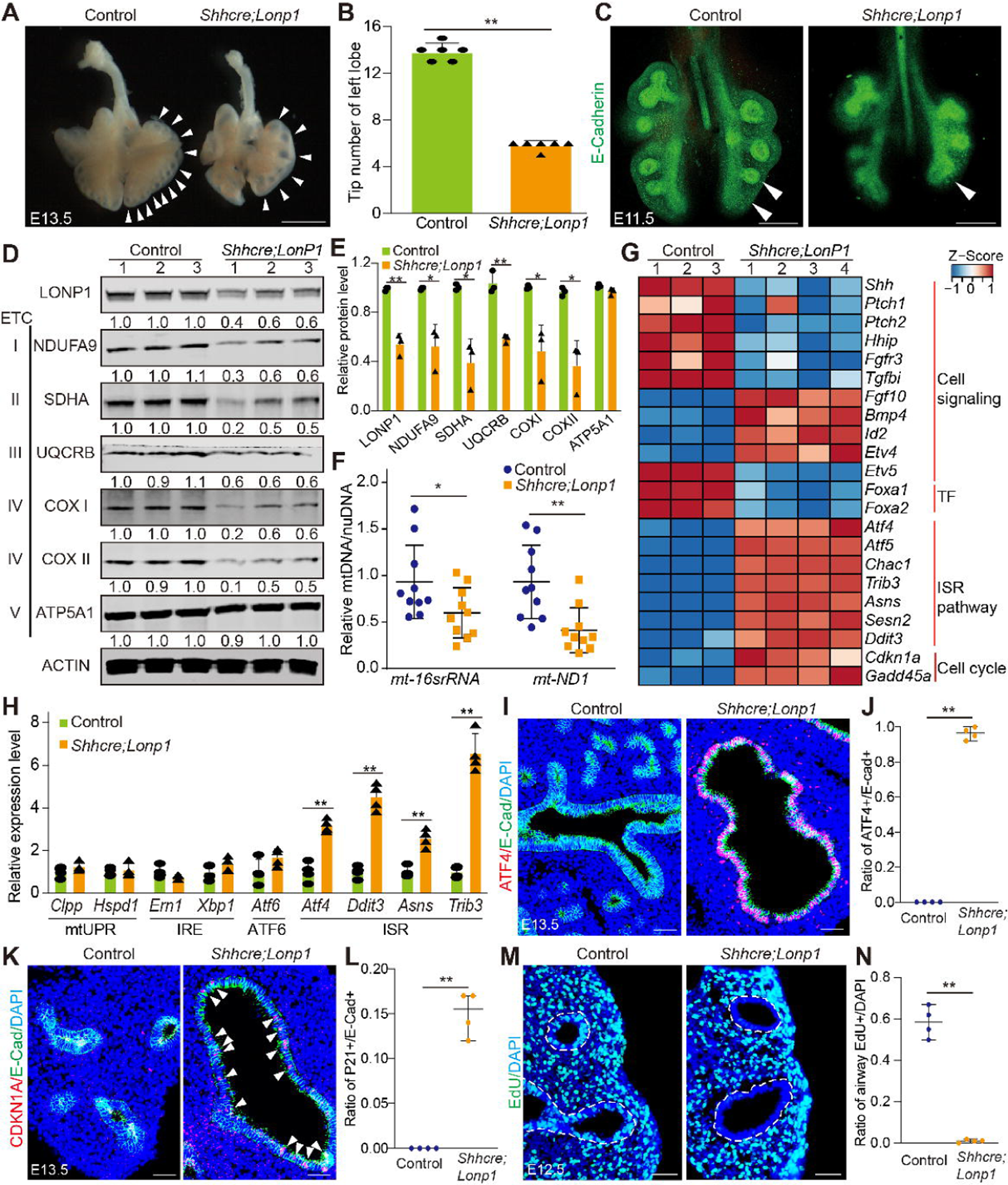
Inactivation of *Lonp1* in the lung epithelium led to branching defects and ectopic activation of the ISR. (A) Lung morphology of *Shhcre;Lonp1* and control at E13.5. Branching tips were denoted by arrowheads. Scale bars, 500 µm. (B) Tip numbers of left lobes at E13.5 (n=6 for each group). (C) Whole mount staining of E-Cadherin at E11.5. Tip number reduction was apparent in the left lobe as denoted by arrowheads. Scale bars, 200 µm. (D-E) Protein levels of representative ETC components in E13.5 lungs as assayed by western blot. Quantifications were shown in (E). n=3 for each group. (F) Quantifications of mitochondria DNA copy number. n=10 for each group. (G) Gene expression levels measured by bulk RNA-seq from E13.5 lungs. (H) qPCR of ER and mitochondria related stress genes at E13.5. n=4 for each group. (I-J) ATF4 immunostaining in E13.5 lungs. E-Cadherin was co-stained to label the epithelium. Quantifications were shown in (J). n=4 for each group. Scale bars, 50 µm. (K-L) CDKN1A immunostaining in E13.5 lungs. E-Cadherin was co-stained to label the epithelium. Nuclear staining of CDKN1A was marked by arrowheads. Quantifications were shown in (L). n=4 for each group. Scale bars, 50 µm. (M-N) Epithelial cell proliferation measured by EdU labeling at E12.5. Epithelium was marked by dashed lines. Quantifications were shown in (N). n=4 for each group. Scale bars, 50 µm.

To determine how loss of *Lonp1* impacts mitochondrial function in lung, we investigated key mitochondria parameters, including protein levels of electron transport chain (ETC) enzyme subunits, the generation of reactive oxygen species (ROS) and total mitochondria numbers. We found that most of the ETC enzymes, including nuclear-encoded subunits NDUFA9 (complex I) and SDHA (complex II), and the mitochondria-encoded subunits UQCRB (complex III), COX I and COX II (complex IV), were significantly decreased in the mutant lung at E13.5, while the nuclear-encoded ATP5A1 (complex V) remain unchanged (Figures 1D and 1E). Mitochondria are the major source of cellular reactive oxygen species (ROS), which is mainly generated from the reduction of oxygen by leaked electrons from ETC during the process of oxidative phosphorylation. To address if ETC defects in *Lonp1* mutants led to changes in ROS production, we quantified mitochondria ROS by using MitoSox staining followed by flow cytometry. We found increased MitoSox staining in the mutant compared to control (Figure S1H). By quantitative PCR, we found a significant reduction of mitochondria DNA copy numbers in the mutant compared to control (Figure 1F). These results together suggested that LONP1 is crucial for mitochondrial homeostasis in the early lung epithelial progenitors.

To comprehensively identify molecular changes in response to *Lonp1* inactivation, we performed bulk RNA-seq in *Shhcre;Lonp1* and control lungs at E13.5. By the cut-off of fold change > 2 and p-value <0.01, 202 up-regulated genes and 143 down-regulated genes were identified in the mutant. Among them, we found that the key epithelial signaling gene, *Shh,* and its mesenchymal targets *Ptch1, Ptch2* and *Hhip*, are decreased in the mutant. Conversely, the SHH repressing mesenchymal signaling gene *Fgf10*, and several of the FGF10 downstream targets including *Bmp4, Id2* and *Etv4* are up-regulated (Figure 1G). Curiously, *Etv5*, another FGF10 downstream target, is downregulated, suggesting complex regulation in the *Lonp1* mutants. These changes were further validated by qRT-PCR (Figure S1M). By *in situ* hybridization, there was a clear expansion of *Fgf10* in the mesenchyme surrounding the tips in the *Lonp1* mutants (Figure S1N). Aside from signaling molecules, we found that two transcription factors, FOXA1 and FOXA2, were both down-regulated in the mutant at both RNA and protein levels (Figures 1G, S1I-S1L). As SHH-FGF feedback loop genes, and *Foxa1*/*Foxa2* are critical for branching morphogenesis (Abler et al., 2009; Herriges et al., 2015; Wan et al., 2005), changes in these genes may contribute to the branching defects observed in *Shhcre;Lonp1* mutants.

A profound signature revealed by the RNA-seq data is the elevated expression of several key genes involved in the integrated stress response (ISR) pathway (Figure 1G), which is further confirmed by qRT-PCR (Figure 1H). This stress pathway activation is highly selective, as neither IRE1 and ATF6 pathways, the other two branches of ER stress response, nor the canonical mitochondrial stress pathway, are affected in the *Lonp1* mutant (Figure 1H). To address if the ISR is activated at the protein level, we performed immunostaining of Activating Transcription Factor 4 (ATF4), the key effector of the ISR (Pakos-Zebrucka et al., 2016). We found a clear increase of ATF4 in the nucleus of epithelial cells of *Lonp1* mutants (Figures 1I and 1J). Previous studies showed that activated ISR could triger either cell death or cell senescence depending on cellular contexts (Pakos-Zebrucka *et al*., 2016). We found no change in apoptosis marker cleaved CASPASE3 staining (Figure S1O), but a clear upregulaton of senescence marker *Cdkn1a* RNA and protein (Figures 1G, 1K-1L), and its direct target *Gadd45a* RNA (Figures 1G). A hallmark of cell senescence is the loss of proliferative capacity. We found that, at E12.5, while control lung epithelium displayed strong proliferative capacity, few to no epithelial cells were proliferative in the mutant lungs as indicated by EdU staining (Figures 1M and 1N). Taken together, these data indicate that impaired mitochondria protein quality control by loss of *Lonp1* has a profound impact on multiple aspects of the embryonic lung epithelium, including increased senescence, decreased proliferation, altered signaling, leading to disrupted growth and branching morphogenesis.

### LONP1 is required for the differentiation and adult homeostasis of airway cells

To investigate if LONP1 is required for cell differentiation, we examined epithelial cell type markers in *Shhcre;Lonp1* lungs compared to control at E18.5. While the AT1 and AT2 markers remain present, there is a near complete loss of both club and ciliated cell markers in the entire mutant epithelium (Figures 2A and 2B). Focusing on trachea where there is normal presence of TRP63+KRT5+ basal cells, while TRP63+ basal cells remain similar, there is a significant increase of TRP63-KRT5+ basal cells extending into the luminal side of the mutant airway (Figures 2C and 2D). These results suggest that LONP1 is required for the differentiation of the airway epithelium.

**Figure 2.**
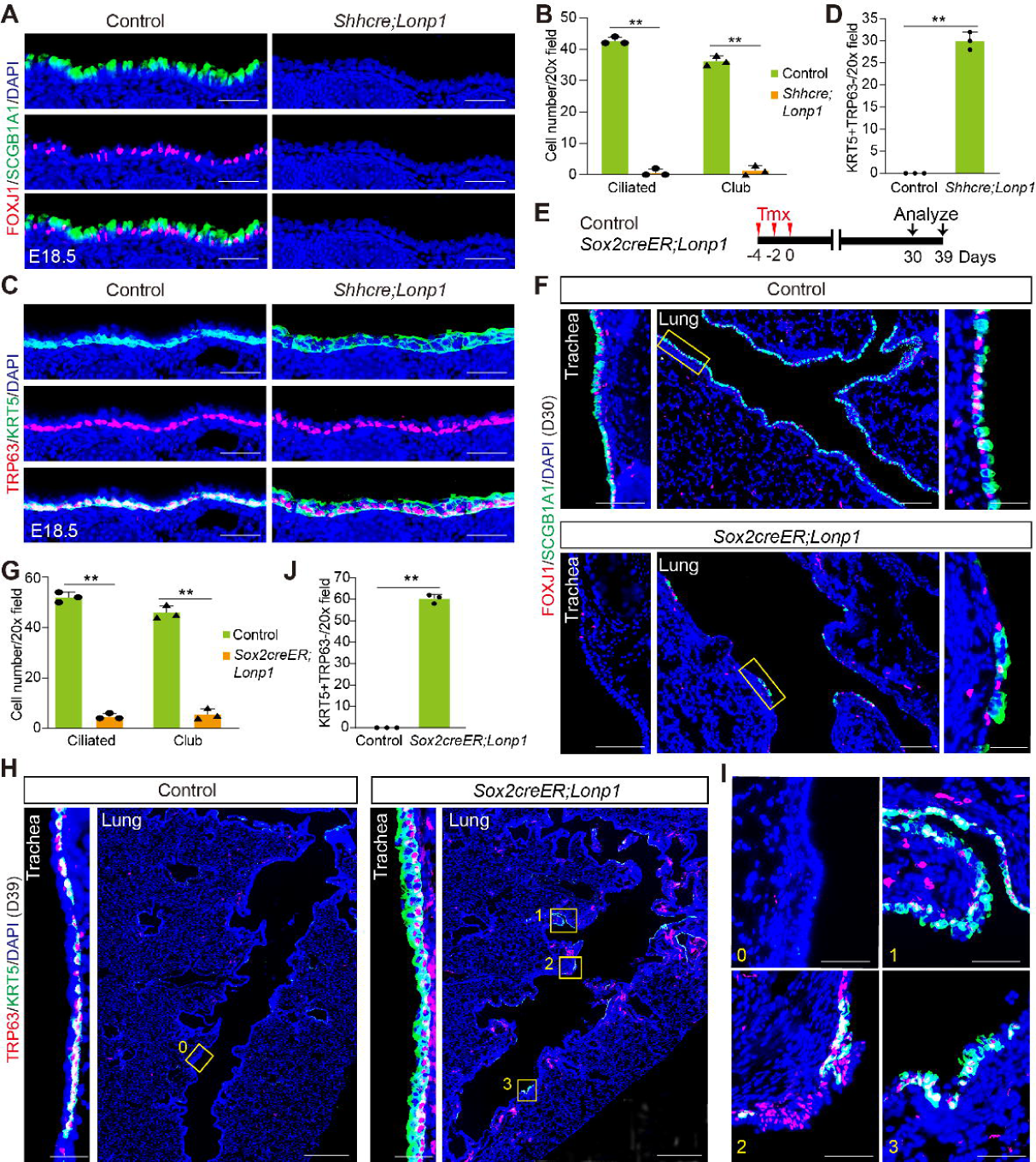
LONP1 is required for differentiation and homeostasis of airway epithelial cells. (A-B) Immunofluorescent labeling of club cells (anti-SCGB1A1) and ciliated cells (anti-FOXJ1) in the trachea of control and *Shhcre;Lonp1* at E18.5. Quantifications were shown in (B). n=3 for each group. Scale bars, 50 µm. (C-D) Immunofluorescent labeling of basal cells by TRP63 and KRT5 at E18.5. Quantifications of luminal staining of KRT5 were shown in (D). n=3 for each group. Scale bars, 50 µm. (E) Schematic of experimental procedure used to adult inactivation of *Lonp1*. (F-G) Immunofluorescent labeling of club and ciliated cells in the trachea and lung of control and *Sox2creER;Lonp1*. The magnified regions of the lung airway were marked by the yellow boxes in the middle image and shown in the right. Quantifications of the trachea staining were shown in (G). n=3 for each group. Scale bars, 50 µm for trachea in the left, 200 µm for lung in the middle and 20 µm for zoomed pictures in the right. (H-J) Immunofluorescent labeling of basal cells in the trachea and lung of control and *Sox2creER;Lonp1*. The magnified regions of the ectopic basal-like cells in the mutant and the corresponding control in (I) were marked by the yellow boxes in (H). Quantifications of the luminal staining of KRT5 in trachea were shown in (J). n=3 for each group. Scale bars, 30 µm for trachea in the left, 500 µm for lung in the middle and 20 µm for zoomed pictures in the right

*Lonp1* is also ubiquitously expressed in adult airway cells based on single cell RNA-seq analysis (Figure S2A) (Montoro et al., 2018), which was further validated by RNAscope co-detections of *Lonp1* RNA with marker proteins for club, ciliated and basal cells (Figure S2B). To determine if *Lonp1* is genetically required for the airway homeostasis, we conditionally inactivated *Lonp1* using *Sox2^creERT2^* (Figure 2E), which is active in all cell types of the adult airway epithelium. Mutant *Sox2^creERT2^;Lonp1^flox/flox^* (hereafter *Sox2creER;Lonp1*) exhibited similar airway abnormalities as *Shhcre;Lonp1*, including a clear reduction of club and ciliated cells (Figures 2F and 2G), and an increase of TRP63-KRT5+ cells in the luminal layer of the trachea (Figures S2C and S2D). Intriguingly, we found that a group of basal-like cells, expressing either TRP63+/KRT5- or TRP63+/KRT5+, ectopically appeared in the intrapulmonary airway of *Sox2creER;Lonp1*, which was rarely seen in the control (Figure S2C). At 39 days post inactivation, these ectopic basal-like cells were found in larger cohorts at diverse regions, some deep in bronchioles with metaplastic appearance (Figures 2H-2J). Taken together, our results demonstrate that LONP1 is essential for the differentiation and adult maintenance of airway epithelial cells.

### LONP1 is uniquely required for the survival of multiciliated cells

Among internal organs, the lung is relatively quiescent at steady state. It is estimated that the terminally differentiated airway epithelial cells have a half-life of ∼6 months in the trachea and ∼17 month in the intrapulmonary airway (Pardo-Saganta et al., 2015; Rawlins and Hogan, 2008). It is therefore intriguing that mutant mice exhibited a severe airway phenotype within one month after *Lonp1* inactivation. In general, cell turnover is largely affected by two processes: a) Replenishment by tissue-resident progenitors, and b) Loss of their progeny. To first determine if *Lonp1* plays a role in airway progenitor self-renewal and differentiation, we assessed *Lonp1*-deficient club and basal cell behaviors under the regenerative condition in *Scgb1a1^creERT2^;Lonp1^flox/flox^* (hereafter *Scgb1a1creER;Lonp1*) (Figure S3A) and *Trp63^creERT2^;Lonp1^flox/flox^;Rosa^tdTomato^* (hereafter *Trp63creER;Lonp1;tdT*) (Figure S3G), respectively. Naphthalene-induced injury led to drastic reduction of club cells in the lung airway, and both club and ciliated cells in the trachea (Figures S3B and S3H). In the regenerative process, *Lonp1*-deficient club cells in the lung maintained a comparable potential for self-renewal as the control ones, evidenced by the nuclear staining of proliferative marker MKI67 (Figures S3C and S3D). At 21 days post injury, previously lost club cells were similarly restored in the mutant airway as in the control (Figures S3E and S3F). In the trachea of *Trp63creER;Lonp1;tdT*, *Lonp1*-deficient basal cells expand to the luminal side of the injured airway at 3 days post injury, and further regressed to the basement membrane while giving rise to the regenerated club and ciliated cells at 9 days post injury (Figures S3I-S3L). We found no significant differences of cell numbers between control and mutant mice at any examined time point (Figures S3M and S3N). As club cells are the direct parents of ciliated cells in the trachea under normal and injured condition (Montoro *et al*., 2018; Rawlins et al., 2009), restoration of ciliated cells in *Trp63creER;Lonp1;tdT* further suggest that club cell differentiation tend to be not affected by loss of *Lonp1*. Collectively, these findings suggest that *Lonp1* is dispensable for the stemness maintenance of the facultative airway progenitors.

We next tested if the airway phenotype of *Lonp1* mutant is caused by the programmed cell death, which is the major form of cell clearance. In *Sox2creER;Lonp1*, we found a clear increase of cleaved Caspase 3 in the mutant epithelium at 14 days post *Lonp1* inactivation (D14), in contrast to its rare presence in the control airway (Figures 3A and 3B). The death signal was noticeable as early as D10, peaked at D22 and remained until the emergence of ectopic basal cells at D30 (Figure 3C). Given cleaved Caspase 3 is not present in every airway cell (Figure 3B, right panel), we wondered if apoptosis preferentially occurred in any specific cell type. To test this, cleaved Caspase 3 was co-stained with major airway cell markers. To our surprise, this death signal was exclusively found in the multiciliated cells, but not in basal stem cells nor club cells (Figures 3D and 3E). To determine if apoptosis was triggered intrinsically by loss of *Lonp1*, we specifically inactivated *Lonp1* in the ciliated cells by generating *Foxj1^creERT2^;Lonp1^flox/flox^* (hereafter *Foxj1creER;Lonp1*) (Figure 3F). Compared to the control in which cleaved Caspase 3 was not detected in any airway cells, the apoptotic signal was evident in the ciliated cells at D18 (Figures 3G and 3H). These data suggest that *Lonp1* plays an intrinsic role in maintaining ciliated cell survival.

**Figure 3.**
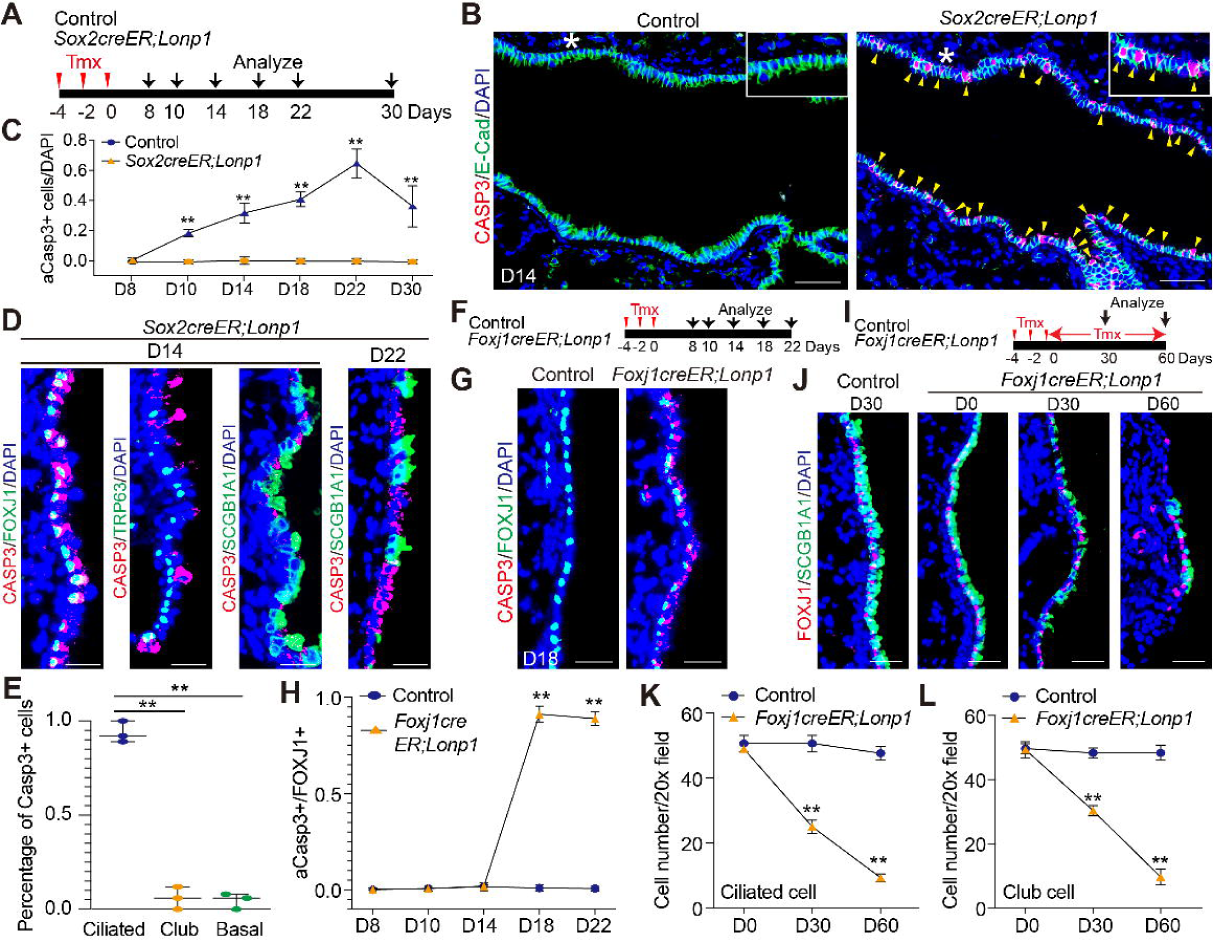
LONP1 is selectively required for the survival of multiciliated cells. (A) Schematic of experimental procedure used to analyze apoptosis in *Sox2creER;Lonp1*. (B) Immunostaining for cleaved Caspase3 in the airway epithelium of control and *Sox2creER;Lonp1* at 14 days post inactivation. Apoptotic cells were denoted by yellow arrows. The magnified regions of the airway were marked by the white stars and shown in the upper right. Scale bars, 50 µm. (C) Ratio of the apoptotic cells in the airway quantified for each time point. (D-E) Co-staining of cleaved Caspase3 with each cell marker for ciliated, basal and club cells at D14 and D22. Quantifications of co-detection at D14 were shown in (E). n=3 for each group. Scale bars, 20 µm. (F) Schematic of experimental procedure used to inactivate *Lonp1* in ciliated cells. (G-H) Staining of cleaved Caspase3 in ciliated cells at D18. Quantifications of co-detection at multiple time points were shown in (H). n=3 for each group. Scale bars, 20 µm. (I) Schematic of experimental procedure used for continuous tamoxifen administration in *Foxj1creER;Lonp1*. (J-L) Immunofluorescent labeling of club and ciliated cells in the lung of control and *Foxj1creER;Lonp1* at D0, 30 and D60 post Lonp1 inactivation. Quantification of ciliated and club cell numbers at denoted time point were shown in (K) and (L), respectively. n=3 for each group. Scale bars, 20 µm.

Previous study showed that one-time ablation of ciliated cells has minor effects on the behaviors of remaining cells in the adult airway (Pardo-Saganta *et al*., 2015). To test if the airway phenotype in *Sox2creER;Lonp1* can be a result from the draining of continuous loss of ciliated cells, ciliated cells alone were subjected to constant cell death by continuous tamoxifen administration to *Foxj1creER;Lonp1* mice (Figure 3I). A significant decrease of both ciliated and club cells was found in the mutant at D30 (Figures 3J-3L). Very few club and ciliated cells were retained in the mutant airways at D60, phenocopying *Sox2creER;Lonp1* (Figures 3J-3L). Interestingly, ectopic basal cells were not found in the intrapulmonary airway of *Foxj1creER;Lonp1* at any time point (data not shown), indicating that the kinetics of luminal cell loss is rate limiting for the differentiation of ectopic basal cells. It is also possible that a synergistic interplay between the *Lonp1*-deficient airway progenitors and their progeny is required for the appearance of these ectopic basal cells.

### LONP1 maintains cell survival by repressing the integrated stress response

We found that the airway epithelium in the mutant lung was not completely denuded from the significant loss of the luminal cells. Aside from the sparsely localized basal-like cells, most of the airway surface were covered by cells with unknown identities (Figures 2F and 2H). To profile the cellular heterogeneity of the mutant airway and dissect the molecular mechanisms underlying the phenotype, we carried out single cell transcriptomic analysis of the lineage-labeled epithelial cells sorted from *Sox2^creERT2^;Lonp1^fl/fl^;Rosa^tdTomato^* and the control lungs (Figure 4A). Totally 27,402 epithelial cells (9,682 in control, 17,720 in mutant) passed our quality control and used for further integration and dimension reduction. 10 clusters were initially identified by unsupervised clustering, and manually annotated by the expression of canonical marker genes (Sun et al., 2022). In addition to the major cell types, a couple of the relatively rare airway cells, including deuterosomal, serous, goblet, BASCs and a group of cycling club cells, were faithfully captured by our protocol (Figure 4B, see Materials and Methods). We found that a population of EpCAM+tdTomato+ aberrant cells was separated from the rest of airway cells by expressing none of the profiled marker genes (Figure 4C). Comparing cells originating from mutant lungs to controls, we found a large population of cells exclusively came from the *Lonp1* mutants (Figure 4D). High-resolution clustering and marker gene identification suggest that this cell population was heterogeneous and consisted of multiple subgroups that shared largely the same marker genes with their counterparts in the control (Figures 4E and 4F). However, these cells were characterized by the decreased expressions of marker genes, and increased expressions of the stress-responsive genes in the ISR pathway, such as *Ddit3, Gadd45a* and *Trib3* (Figure 4F). Consistent with the results from immunostaining, relative proportions of control and mutant cells in each cell type demonstrated that cell numbers of club and ciliated cells were greatly decreased in the mutant airway, while basal cell numbers were increased there (Figure 4G). In comparison, rare cell types such as deuterosomal and goblet cells, as well as cells with stressed signature, were more enriched in the *Lonp1* mutants (Figure 4G).

**Figure 4.**
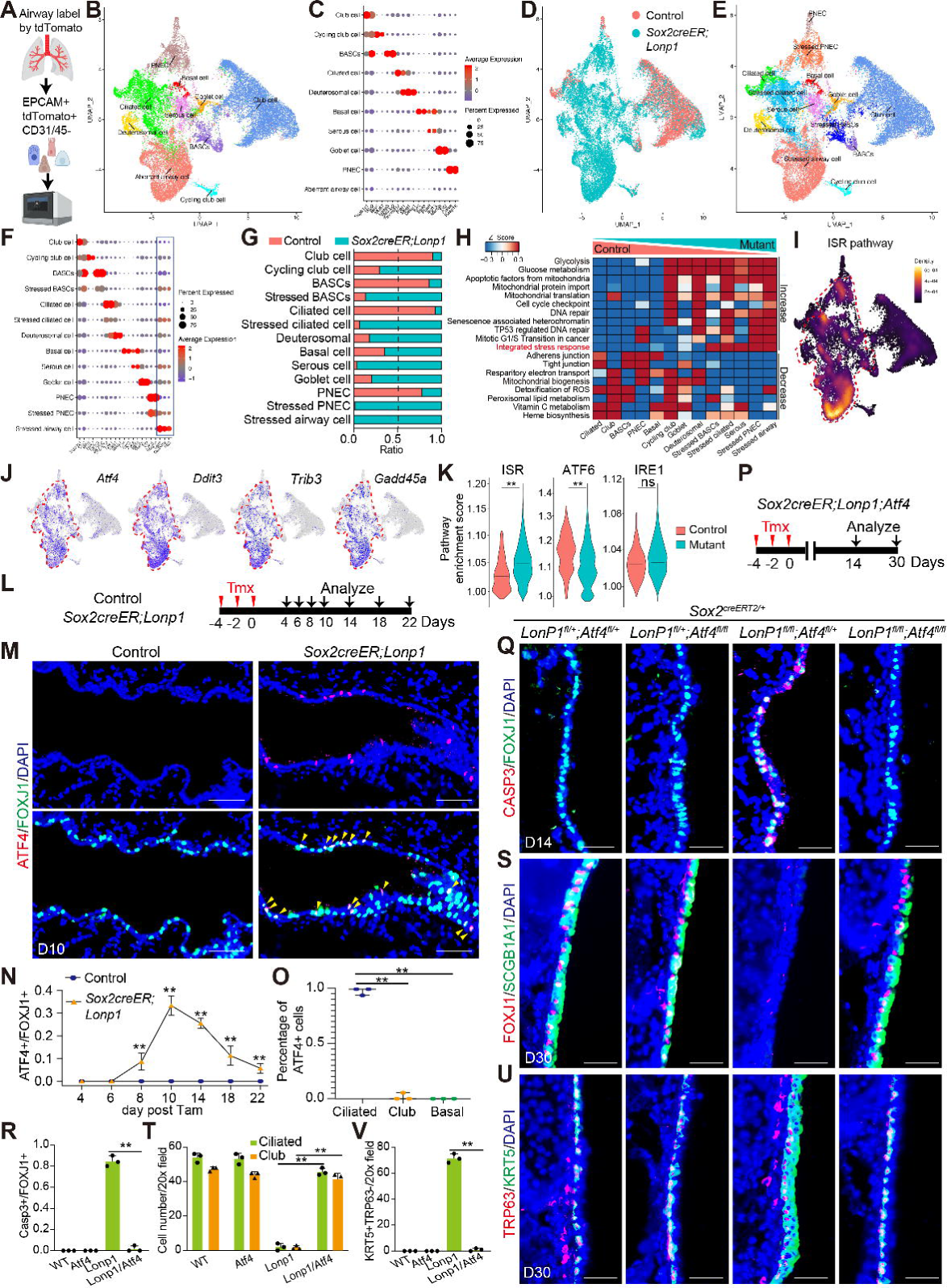
Ectopic activation of the ISR pathway in ciliated cells led to apoptosis. (A) Schematic of experimental procedure used to isolate airway epithelial cells for scRNA-seq. (B) UMAP embedding of integrated cells from control and *Sox2creER;Lonp1*. (C) Expression signatures of marker genes in each cell type. (D) UMAP embedding of integrated cells colored by control and *Sox2creER;Lonp1*. (E) UMAP embedding of integrated cells with higher resolution. (F) Expression signatures of marker genes in each defined cell type. (G) Ratio of normalized cell numbers from control and *Lonp1* mutant in each cell type. (H) Differentially enriched pathways between control and mutant airway cells analyzed by gene set enrichment analysis. Cell types were ordered horizontally by the relative proportions of control and mutant cell numbers, as denoted by the right triangles on the top. (I) Single cell enrichment scores of the ISR pathways projected onto UMAP. Cells exclusively identified in the mutant airway were marked by red dashed lines. (J) Gene expression features of representative factors in the ISR pathway. Cells exclusively identified in the mutant airway were marked by red dashed lines. (K) Distribution of enrichment scores for three branches of ER stress pathway in control and mutant cells. Wilcox test was used for statistical assay. (L) Schematic of experimental procedure used to analyze the ISR activation in *Sox2creER;Lonp1*. (M) Co-immunostaining of ATF4 with FOXJ1 in the airway epithelium of control and *Sox2creER;Lonp1* at 10 days post inactivation. Scale bars, 50 µm. (N) Ratio of the ATF4+ ciliated cells in the airway quantified for each time point. (O)Quantifications of ATF4 co-staining with ciliated, club and basal cell markers at 10 days post inactivation. n=3 for each group. (O) Schematic of experimental procedure used to assay airway phenotypes in *Sox2creER;Lonp1;Atf4*. (Q-R) Immunofluorescent labeling of apoptotic signals on top of FOXJ1 in the trachea of control and relative mutants at 14 days post inactivation. Quantifications were shown in (R). n=3 for each group. Scale bars, 20 µm. (S-T) Immunofluorescent labeling of club and ciliated cells in the trachea of control and relative mutants at 30 days post inactivation. Quantifications were shown in (T). n=3 for each group. Scale bars, 20 µm. (U-V) Immunofluorescent labeling of basal cells in the trachea of control and relative mutants at 30 days post inactivation. Quantifications were shown in (V). n=3 for each group. Scale bars, 20 µm.

To comprehensively profile the molecular changes within the *Lonp1*-deficient airway cells, we performed the pathway enrichment analysis at single cell level. By comparing gene set enrichment scores between control and mutant cells, we found that some mitochondria-associated metabolic processes, such as vitamin C and heme biosynthesis, were found reduced in the mutant-enriched cell types (Figure 4H). Instead, glycolysis, cell cycle checkpoint, P53-mediated DNA repair and integrated stress response pathways were significantly elevated (Figure 4H). Projection of enrichment score of the ISR on UMAP (Figure 4I), as well as the RNA expression features of the ISR hallmarks (Figure 4J), revealed that the ISR is broadly activated in the mutant airway epithelium. Consistent with our findings in embryonic *Lonp1* mutant, other two branches of ER stress were not elevated in the stressed cells (Figure 4K).

We next assessed the ISR activation at the protein level by immunostaining for ATF4 in *Sox2creER;Lonp1* and its corresponding control at multiple time points (Figure 4L). We found that, in the mutant airway, ATF4 protein was initially detected in the nucleus of ciliated cells at D8 (Figure 4N), and became most abundant at D10 when about 30% of ciliated cells possessed nuclear ATF4 (Figures 4M and 4N). Consistent with the previous Caspase 3 staining, neither club nor basal cells were found possess nuclear ATF4 (Figure 4O). The discrepancy between RNA and protein signatures of ATF4 in the *Lonp1* mutant airway at adult stage indicate the potential regulation of ATF4 at the post-transcriptional level. We speculate that the survival of ciliated cells is maintained by the cell autonomous suppression of the ISR by LONP1. To test our hypothesis *in vivo*, we sought to inactivate *Atf4* in the *Lonp1* mutant background by generating *Sox2^creERT2^;Lonp1^flox/flox^;Atf4^flox/flox^ (hereafter Sox2creER;Lonp1;Atf4)* (Figure 4P). Assay of cell death revealed that apoptotic process induced by loss of *Lonp1* is largely blocked by *Atf4* inactivation (Figures 4Q and 4R). Importantly, the loss of club and ciliated cells in the *Lonp1* single mutant were restored by the additional inactivation of *Atf4* (Figures 4S and 4T). The increase of basal cells in the trachea of *Lonp1* single mutants was also attenuated in the *Sox2creER;Lonp1;Atf4* (Figures 4U and 4V).

It had been shown that ATF4 mediates cell apoptosis mainly through the downstream death factor DDIT3 (Oyadomari and Mori, 2004). We next inactivated *Ddit3* in the *Lonp1* mutant background by generating *Sox2^creERT2^;Lonp1^flox/flox^;Ddit3^flox/flox^ (hereafter Sox2creER;Lonp1;Ddit3)* (Figure S4A). The reduction of both club and ciliated cells was largely rescued in the *Sox2creER;Lonp1;Ddit3* (Figures S4B and S4C), as well as increase of basal cells (Figures S4D and S4E). Noticeably, it is unlikely that these genetic rescues were the consequence of inefficient Cre-mediated recombination, since similar levels of reduction in *Lonp1* RNA levels were achieved between single and double mutants (Figures S4F and S4G). These findings together suggest that LONP1-repressed ISR pathway is the direct cause of ciliated cell death, leading to a cascade of airway phenotypes including the loss of club cells and basal cell increase.

### LONP1 renders migratory capacity of resident airway progenitors upon virus-induced lung injury

It has been shown that dysplastic KRT5+ basal cells are induced in the airway post influenza infection, and migrate into the injured alveoli. To investigate if LONP1 plays a role in mediating such dysplastic response, we inactivated *Lonp1* in the airway stem cells while performing the lineage tracing by using *Sox2^creERT2^;Lonp1^fl/fl^;Rosa^tdTomato^* (hereafter *Sox2creER;Lonp1;tdT*) (Ray et al., 2016; Xi et al., 2017). To prevent the influence of baseline phenotypes associated with long-term loss of *Lonp1*, both the control and mutants were subjected to intranasal inoculation of the mouse-adapted influenza (H1N1/PR8) 4 days after the last injection of tamoxifen, and lungs were analyzed 14 days post infection (Figure 5A). Unlike in the control that the airway derived KRT5+ basal cells migrated into the damaged alveolar region and formed “KRT5+ pods”, they were rarely seen in the damaged alveoli of *Sox2creER;Lonp1;tdT* (Figures 5B-5D). This phenotype is unlikely to be a result of decreased susceptibility to viral infection due to *Lonp1* inactivation, since both body weight loss (Figure S5A) and viral load (Figure S5B) in the mutants exhibited evidence of an even more severe infection than in control mice. It has been shown that the appearance, and the subsequent expansion, of KRT5+ cells are largely restricted in the airway before 11 days post infection, and these cells migrate into the injured alveoli afterwards (Ray *et al*., 2016; Vaughan *et al*., 2015). We found that KRT5+ cells in the *Lonp1* mutant were still capable of expansion in the airway, at a similar level as control (Figures 5C and 5E). Collectively, these *in vivo* data suggest that LONP1 is required for the alveolar migration of airway KRT5+ basal cells post influenza infection.

**Figure 5.**
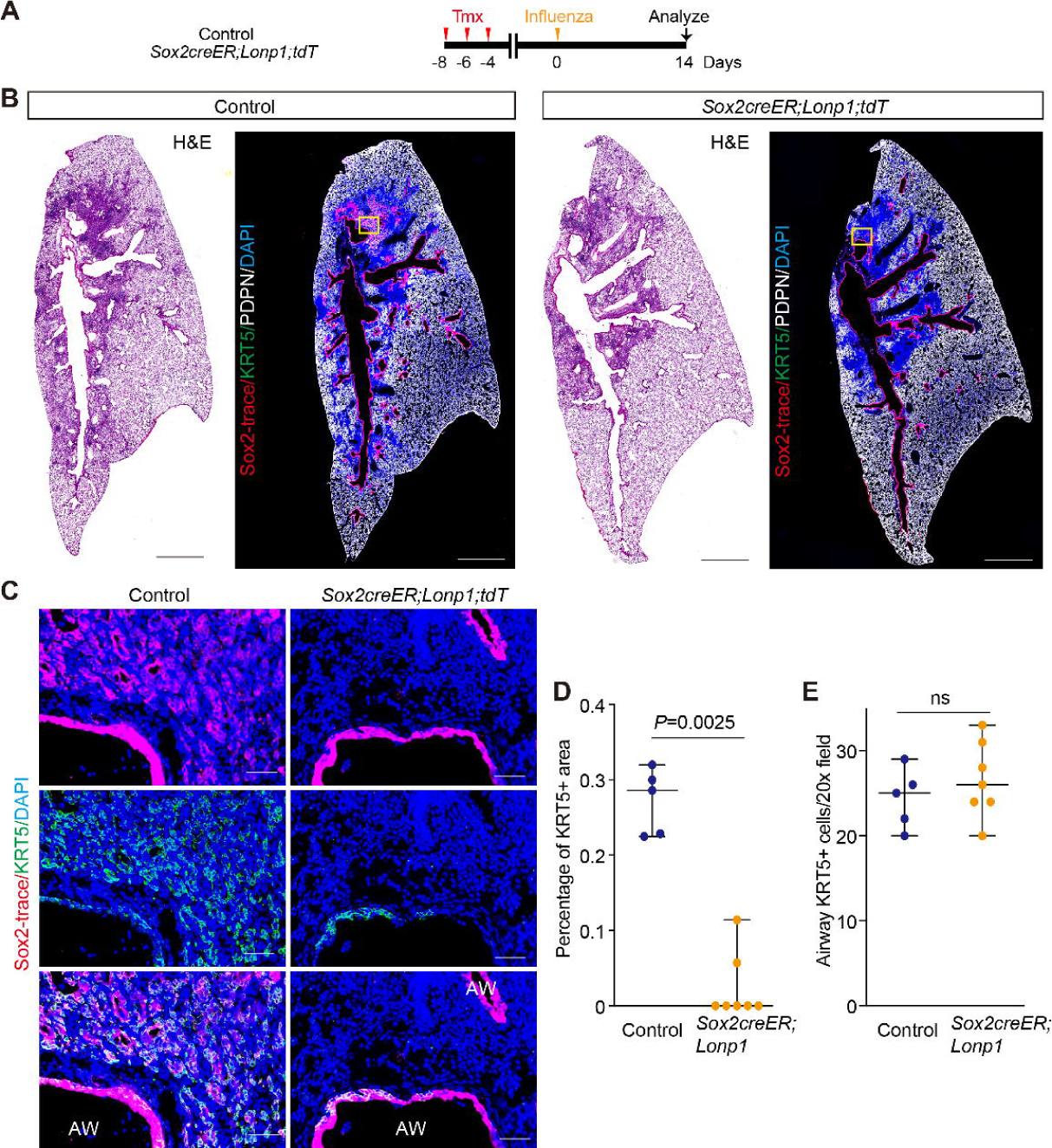
LONP1 is required for the alveolar migration of ectopic KRT5+ basal cells post influenza infection. (A) Schematic of experimental procedure used to perform influenza infection to *Sox2creER;Lonp1;tdT* and the corresponding control. (B) Histological morphology of virus injured lungs measured by H&E staining, and the lineage tracing of KRT5+ airway stem cells in the adjacent sections on the right. Severe viral injury led to drastic loss of AT1 cells, which was evidenced by the absence of PDPN staining. Scale bars, 1 mm. (C) Magnified regions marked by yellow boxes in (B), which showed the stalled KRT5+ basal cells in the airway of *Sox2creER;Lonp1;tdT.* Scale bars, 50 µm. (D) Percentage of viral damaged areas covered by KRT5+ basal cells in controls and *Lonp1* mutants. n>=5 for each group. (E) Number of KRT5+ basal cells in the injured airways of controls and *Lonp1* mutants. n>=5 for each group.

### LONP1 promotes formation of alveolar KRT5+ pods through repression of the ISR

A few signaling pathways involved in maintaining the stemness of progenitors during lung development are reutilized as part of the injury response to promote cellular plasticity. We next assessed if LONP1 controls the formation of KRT5+ pods through repressing the ISR pathway. Immunostaining revealed that ATF4 is highly expressed in the nucleus of KRT5+ cells in the airway of *Sox2creER;Lonp1;tdT*, but not in the control mice, post influenza infection (Figures 6A and 6B). This suggests the derepressing of the ISR in the stalled basal cells of the mutant airway. To determine the downstream cellular outcome, we analyzed markers of senescence and apoptosis. While no evidence of apoptosis was observed (Figure 6C), increased staining of cell cycle arrest factor CDKN1A was found in the basal cells of the mutant (Figures 6D and 6E). To determine if the activation of the ISR is the direct cause of the basal cell stalling, we carried out influenza infection in aforementioned *Sox2creER;Lonp1;Atf4* double mutants, aside from the *Lonp1* single mutants (Figure 6F). Intriguingly, KRT5+ pods were found reappear in the damaged alveoli in the double mutants (Figures 6G-6I). These findings demonstrate that the migratory capacity of KRT5+ basal cells post influenza injury rely on the cell autonomous suppression of the ISR by LONP1.

**Figure 6.**
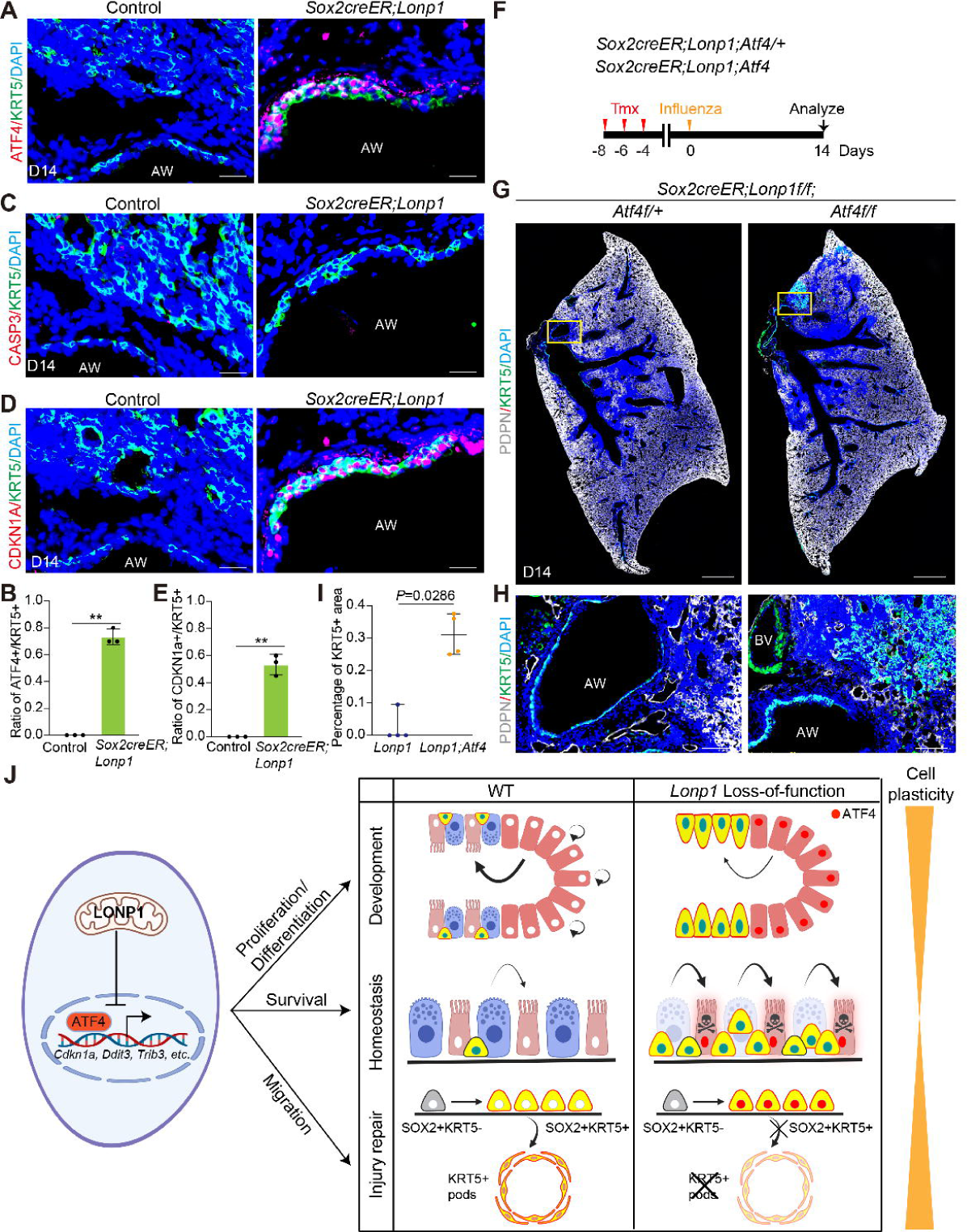
LONP1 maintains the migratory capacity of ectopic KRT5+ basal cells by repressing the ISR pathway. (A-B) Immunofluorescent labeling of ATF4 with ectopic KRT5+ basal cells in the airway of control and *Sox2creER;Lonp1* at 14 days post influenza infection. Quantifications were shown in (B). n=3 for each group. Scale bars, 20 µm. (C) Cleaved Caspase3 staining was not found neither in control nor in mutant airways. Scale bars, 20 µm. (D-E) Immunofluorescent labeling of CDKN1A with ectopic KRT5+ basal cells in control and *Sox2creER;Lonp1* at 14 days post influenza infection. Quantifications were shown in (E). n=3 for each group. Scale bars, 20 µm. (F) Schematic of experimental procedure used to perform influenza infection to *Sox2creER;Lonp1;Atf4* and *Sox2creER;Lonp1;Atf4/+*. (G-I) Immunofluorescent labeling of KRT5 at 14 days post influenza infection. Magnified regions marked by yellow boxes in (G) were shown in (H), percentage of KRT5+ areas in the damaged alveoli were quantified in (I). n=4 for each group. Scale bars, 1 mm in (G) and 100 µm in (H). (J) LONP1-mediated mitochondria protein quality control plays an array of essential roles in regulating airway epithelial cell proliferation, differentiation, survival and migration in a context dependent manner. Cartoons were created with BioRender.com.

## DISCUSSION

In this study, we provide evidence from *in vivo* experiments demonstrating that mitochondria protein quality control, which is modulated by LONP1, dynamically regulates cell behaviors of airway epithelium through repression of the integrated stress response (Figure 6J). For airway progenitors with increased plasticity during early lung development and during regenerative processes, LONP1 functions to protect cells against cellular senescence, promote proliferation and differentiation of naïve epithelial progenitors, and maintain migratory capacity of basal-like cells post influenza-induced injury. In contrast, during adult airway homeostasis, LONP1 is selectively required for the survival of terminally differentiated ciliated cells. Genetic rescues by *Atf4* deficiency suggest that the ISR is the primary pathway downstream of *Lonp1* loss of function which drives these phenotypes.

Our findings here raise an interesting question about how distinct cell types respond in different ways to the same mitochondria abnormality. It has been shown that cells with a high energy demand tend to have greater numbers of mitochondria (Kukat et al., 2011), and the intracellular distribution of mitochondria is responsible for specific cell function and activity (Frederick and Shaw, 2007). The metabolic state of the cell also significantly changes the cellular demand for energy and the substrates generated from mitochondria (Mishra and Chan, 2016). These features allow for the differing susceptibility of cell types to the mitochondrial perturbations. The high-frequency beating of cilia on multiciliated cells in the airway is crucial for the clearance of the mucus-trapped particles from the respiratory tract, which relies on the availability of massive ATPs produced by mitochondria that are localized adjacent to the basal bodies of cilia (Kikkawa, 2013). Our findings that LONP1 selectively maintains the survival of ciliated cells implies that these cells are sensitive to mitochondrial perturbation probably due to their high energy requirements. This, in turn, provides long-term adaptive benefits to efficiently turnover and replenish terminally differentiated ciliated cells to maintain airway epithelial health throughout adulthood.

*In vitro* studies indicate that ATF4 activation normally contributes to stress relief and cell survival, but can upregulate apoptotic cascades in response to persistent stimuli (Wortel et al., 2017). Here we show for the first time, to our knowledge, that organ-wide activation of the ISR led to either pro-survival senescence of progenitors, or apoptosis of their terminally differentiated progeny. How cell survival and apoptosis are controlled downstream of ISR activation is not well understood. It has been shown that ATF4 heterodimerization with different binding partners is associated with the switch between pro-survival and pro-apoptotic signaling (Ameri and Harris, 2008; Han et al., 2013). Post-translational modifications of ATF4 protein are identified to affect its stability and transcriptional activity (Lassot et al., 2005; Lassot et al., 2001; Yuniati et al., 2016). In addition, multiple chromatin modifications on target genes are required for transcriptional regulation by ATF4 (Shan et al., 2012; Zhao et al., 2016). We speculate that these context dependent mechanisms, either alone or in combination, serve as key factors that contribute to the distinct cell behaviors of airway epithelium in *Lonp1* mutants.

Chemical agents are widely used to model acute airway injury in order to study the turnover of epithelial cells, which are either non-cell type specific (polidocanol, SO2) or only functional to subsets of club cells expressing cytochrome P450 (naphthalene). A genetic strategy to ablate ciliated cells by DTA found minimal activation of airway progenitors (Pardo-Saganta *et al*., 2015), while club cell ablation led to extensive proliferation of the remaining progenitor cells (Perl et al., 2011). It is noteworthy that injuries modulated by these approaches are a one-time event, in which no further damages are retained in the replenished cells unless iteratively administrated. How airway stem cells respond to the constant loss of their terminally differentiated progeny, and the global consequences it may cause, remain largely unknown. Given its selective requirement for the survival of ciliated cells, LONP1 provides a unique entry point to address this question. Through universal inactivation of *Lonp1* in entire airway epithelium, or continuous exposure of *Foxj1creER;Lonp1* to tamoxifen, we show that constant loss of ciliated cells led to a significant decrease of their direct progenitor, club cells. As a wound response in many settings of acute lung injury, basal stem cells proliferate and repopulate the denuded airway. Our single cell transcriptomic analysis further demonstrates that some rare cell types, such as deuterosomal and goblet cells, exhibit increased cell numbers in the remodeled airway of *Lonp1* mutants. Aspects of these phenotypes recapitulate common morphological changes of the airway found in COPD and asthma, in which pseudostratified epithelium is replaced with squamous basal cells, and a reduction in club and ciliated cell numbers, aside from an increase of goblet cells, are frequently reported (Gamez et al., 2015; Gohy et al., 2019; Pilette et al., 2001). Our findings here raise the possibility that disrupted mitochondria protein quality control may serve as an additional contributor to these complex airway diseases.

Following influenza-induced injury, the ectopic bronchiolization of damaged alveoli by airway-resident basal-like cells undergoes two major steps that are characterized in a spatially and temporally specific manner: a) Activation and expansion within the injured airway (D0-D11 post infection), and b) migration from the airway and repopulation of the damaged alveoli (D11-D14 post infection) (Ray *et al*., 2016; Vaughan *et al*., 2015). The differentiation and migratory activity of basal-like cells are non-reversible and represent the permanent loss of gas-exchange surface, since dysplastic basal cells in the alveoli rarely differentiate into alveolar cell types and persist as an “epithelial scar” long after injury (Weiner et al., 2022). Though transcription factors, including HIF1a and TRP63, have been shown to play critical roles in rewiring the transcriptional programs that potentiate the migration (Weiner *et al*., 2022; Xi *et al*., 2017), cell-autonomous signaling pathways outside the nucleus that modulate the migratory process remain to be elucidated. Here we discovered that the cell autonomous suppression of the ISR pathway by mitochondria LONP1 protects resident airway stem cells against cell cycle arrest, and maintains their migratory capacity. This auto-inhibitory pathway mediated by mitochondria protein quality control machinery likely functions as a rheostat to fine-tune the dysplastic response of basal-like cells by integrating environmental cues from the injury setting.

In summary, our findings here reveal that LONP1-mediated mitochondria protein quality control plays an array of essential roles in regulating airway epithelial cell proliferation, differentiation, survival and migration in a context dependent manner. This study provides novel insights into the mitochondria-mediated modulation of airway cell fates. Harnessing of key factors in the related pathways allows for the development of novel therapeutical interventions for many airway diseases.

## METHODS

### Mice

*Shh^cre^*, *Tbx4rtTA*, *tetOcre* and *Lonp1^flox^* alleles and transgenic lines have been described previously (Harfe et al., 2004; Qiao *et al*., 2021; Zhang et al., 2013). *Sox2creERT2, Trp63creERT2, Krt5creERT2, Scgb1a1creERT2, Foxj1creERT2, R26RLSL-tdTomato* and *Ddit3^flox^* mice were purchased from JAX. *Atf4^flox^* was a kind gift from Dr. Michael Karin at UCSD with permission from Dr. Christopher Adams at University of Iowa. Embryos used in this study were harvested from time-mated mice, counting noon the day when the vaginal plug was found as E0.5. All mice were in B6 background or have been back crossed to B6 background for at least 3 generations. Littermates were used as controls in all experiments. Tamoxifen dissolved in corn oil was administrated intraperitoneally at the dose of 50 mg/kg. All mice were housed in facilities accredited by American Association for Accreditation of Laboratory Animal Care (AAALAC) at University of California San Diego. All animal husbandry and experiments were approved by Institutional Animal Care and Use Committee (IACUC).

### Tissue preparation and immunofluorescent staining

Tracheas and lungs from embryos or adults were fixed in 4% paraformaldehyde (Electron Microscopy Sciences) diluted in PBS for overnight at 4°C. Samples were embedded in either paraffin or OCT (Electron Microscopy Sciences) for sectioning. Antigen retrieval was performed before serum-mediated blocking by using high-PH retrieval buffer (10 mM Tris, 1 mM EDTA, pH 9.0). Primary antibodies with final concentrations used for immunofluorescence staining: rabbit anti-SCGB1A1 polyclonal antibody [5 mg/ml] (WRAB-3950, Seven Hills Bioreagents), mouse anti-FOXJ1 monoclonal antibody [8 mg/ml] (14-9965-80, eBioscience), mouse anti-TRP63 monoclonal antibody [8 mg/ml] (CM163a, Biocare Medical), rabbit anti-KRT5 polyclonal antibody [5 mg/ml] (905501, Biolegend), rabbit anti-SOX2 polyclonal antibody [5 mg/ml] (NB110-37235, Novus Biologicals), mouse anti-SOX9 monoclonal antibody [5 mg/ml] (14-9765-82, eBioscience), mouse anti-E-Cadherin monoclonal antibody [5 mg/ml] (610181, BD Transduction Laboratories), rabbit anti-FOXA1 monoclonal antibody [5 mg/ml] (ab173287, Abcam), rabbit anti-FOXA2 monoclonal antibody [5 mg/ml] (ab108422, Abcam), rabbit anti-Cleaved Caspase-3 polyclonal antibody [5 mg/ml] (9661, CST), rabbit anti-ATF4 monoclonal antibody [5 mg/ml] (11815, CST), rabbit anti-CDKN1A monoclonal antibody [5 mg/ml] (ZRB1141, Sigma), syrian hamster anti-PDPN polyclonal antibody [5 mg/ml] (Developmental Studies Hybridoma Bank). The following secondary antibodies were used with final concentration: Cy3-conjugated goat anti-mouse IgG [2 mg/ml], Cy3-conjugated goat anti-rabbit IgG [2 mg/ml], AF488-conjugated goat anti-mouse IgG [2 mg/ml], AF488-conjugated goat anti-rabbit IgG [2 mg/ml]. AF647-conjugated goat anti-syrian hamster IgG [2 mg/ml]. All images were acquired on ZEISS AxioImager 2. 20X IF images were used to quantify cells labeled by specific markers. For each sample, at least 3 sections per mouse, and 3 mice per genotype were analyzed.

### EdU analysis for cell proliferation

Click-it EdU Cell Proliferation Kit (C10337, Invitrogen) was used to measure the proliferation signal. For EdU analysis in embryos, 1 ml of 400 mM EdU solution (diluted in PBS, Invitrogen) was intraperitoneally injected into pregnant females. Lungs were harvested 2 hours post injection. Samples were fixed in 4% PFA overnight at 4°C before OCT embedding.

### RNA in situ hybridization and RNAscope

Lungs from embryos were dissected in cold PBS, fixed in 4% paraformaldehyde overnight at 4°C, and then dehydrated to 100% MeOH. Whole-mount in situ hybridization was performed according to established protocol (Abler et al., 2011). For RNAscope, multiplex Fluorescent v2 Assay Kit (ACD) was used to detect Lonp1 RNA in the adult trachea. Fresh-frozen sample were prepared and processed following the RNAscope protocol. *Lonp1* RNA probe stained with the Opal 570 fluorophore were then counter stained with DAPI and FOXJ1/SCGB1A1/TRP63 antibodies as described above.

### Mitochondria DNA copy number analysis

Analysis of the mitochondria DNA copy number is performed according to published protocol (Quiros et al., 2017).Briefly, embryos at E13.5 were harvested and subjected to total DNA extraction. Two pairs of qPCR primers specific to mitochondria DNA (mtDNA) were used for quantification by qPCR: a) 16S rRNA, FWD: 5’-CCGCAAGGGAAAGATGAAAGAC-3’, REV: 5’-TCGTTTGGTTTCGGGGTTTC-3’ b) ND1, FWD: 5’-CTAGCAGAAACAAACCGGGC-3’, REV: 5’-CCGGCTGCGTATTCTACGTT-3’. Nuclear DNA (nDNA) specific gene HK2 (FWD: 5’-GCCAGCCTCTCCTGATTTTAGTGT-3’, REV: 5’-GGGAACACAAAAGACCTCTTCTGG-3’) was used for normalization. The final ratio of mtDNA/nDNA was used as the readout for mtDNA copy number variation.

### Mitochondria ROS measurement by MitoSOX

The specific detection and measurement of mitochondria ROS in embryonic epithelium was conducted by using MitoSOX Red mitochondrial superoxide indicator (Invitrogen). Embryonic lungs at E13.5 were harvested and disassociated in the lysis buffer (PRMI 1640 (Thermo Scientific) with 10% FBS, 1mM HEPES (Life Technology), 1mM MgCl2 (Life Technology), 1mM CaCl2 (Sigma-Aldrich), 0.525mg/ml collagenase D (Roche), 5 unit/ml Dispase (Stemcell Technologies) and 0.05 mg/ml DNase I (Roche)). Incubation of the lysate on the rotator at 150 rpm at 37°C for 15 min, followed by the removal of red blood cells by RBC lysis buffer (Biolegend). Remaining cells were resuspended and blocked by FcBlock antibody (5 mg/ml, BD) for 10 min at RT. Then following antibodies were added to label the corresponding cells: CD326-APC (118213, Biolegend) for epithelial cells, CD31-AF700 (102444, Biolegend) for endothelial cells, CD45-BV510 (103137, Biolegend) for immune cells, Ghost dye red 780 for cell viability (13-0865, TONBO) and the MitoSOX. After 20 min incubation at RT, cells were fixed with the BD stabilizing fixative and kept in 4°C until FACS. FlowJo software (BD) was used to analyze the acquired data.

### Quantitative PCR (qPCR)

Total RNA from embryonic and adult lungs were extracted by using Trizol (Invitrogen) and RNeasy Micro RNA extraction kit (Qiagen). Reverse transcription was then carried out to obtain corresponding cDNA using iScript Select cDNA Synthesis Kit (Bio-Rad). qPCR master mix was prepared using SYBR Green reagents (Bio-Rad) and amplification signals were obtained by CFX ConnectTM system (Bio-Rad). At least three biological replicates were assayed for each gene unless otherwise notated. Primers used for qPCR analysis are listed in Supplemental Table 1.

### Western blot analysis

Embryonic lungs at E13.5 were collected in RIPA Lysis and Extraction Buffer (Thermo) supplemented with cOmplete Protease Inhibitor Cocktail tablets (Roche). Samples were homogenized in a QIAGEN TissueLyser II and proteins were purified from the supernatant after high-speed centrifuge. Protein concentration were measured by PierceTM BCA Protein Assay kit (Thermo Scientific). 10 µg protein sample was loaded in each well and ran on a 4%–12% SDS-PAGE gel (Invitrogen), then transferred to the nitrocellulose membrane (0.2 µm, Bio-Rad) by the Trans-Blot Turbo Transfer System (Bio-Rad). Then the membranes were blocked in TBST (0.1% Tween-20) with 5% BSA for 1.5 hours at room temperature (RT) before incubated in primary antibodies overnight at 4°C. After 3 rounds of wash with TBST, membranes were incubated in the secondary antibody for 1 hour at RT. Images were acquired by Odyssey DLx Imager (LI-COR) and quantified by ImageJ. The following primary antibodies were used: LONP1 (28020, CST, 1:1000), NDUFA9 (459100, Invitrogen, 1:1000), SDHA (5839S, CST, 1:1000), UQCRB (10756-1-AP, Proteintech, 1:1000), COX I (ab14705, Abcam, 1:500), COX II (55070-1-AP, Proteintech, 1:1000), ATP5A1 (14676-1-AP, Proteintech, 1:500) and beta-Actin (Novus Biologicals, NB600501, 1:5000). Secondary antibodies: donkey anti-Mouse IgG IRDye 680RD (LI-COR, 926-68072, 1:10000) and donkey anti-Rabbit IgG IRDye 800CW (LI-COR, 926-32213, 1:10000). Three biological replicates were analyzed for each sample.

### Bulk RNA-seq and data analysis

Total RNA from embryonic lungs at E13.5 were extracted as previously described. cDNA libraries were constructed by using Illumina TruSeq RNA Library Prep Kit V2 (Illumina) and sequenced on the HiSeq4000 platform (Illumina) at the Institute for Genomic Medicine (IGM) at UCSD. FASTQ files were aligned to the mouse reference genome (mm10) by using Bowtie2 (Langmead and Salzberg, 2012) with default settings. Differential gene expression analysis was performed using Cufflinks (Trapnell et al., 2010). Heatmaps were generated by ggplot2 (version 3.3.2).

### Tissue dissociation and sorting of epithelial cells

*Sox2creER;Lonp1;tdTomato* and the corresponding control mice at 39 days post *Lonp1* inactivation were used for cell disassociation followed by scRNA-seq. Briefly, to remove the circulating blood cells, every mouse was transcardially perfused with 12 ml of cold DPBS (Life Technology) before harvest. Lungs were then inflated intratracheally with cold lysis buffer (recipe referred to the mitoSOX method). Both trachea and lungs were chopped into the lysis buffer and mechanically homogenized by GentleMACS dissociator (Miltenyi Biotec). Tissues were further incubated in the lysis buffer on the rotator (150 rpm) for 30 min at 37°C, then filtered through a 70 µm MACS SmartStrainer. Red blood cells were removed by ACK lysis buffer (Gibco). Cells were further collected by centrifuge at 1500 rpm at 4°C for 5 min (acceleration/deceleration=7), counted with hemocytometer and then diluted to ∼1*10^6^ cells per ml. After blocking with FcBlock antibody (5 mg/ml, BD) for 10 min at RT, endothelial cells and immune cells were labeled with biotinylated CD31 (13-0311-82, eBioscience) and CD45 (13-0451-82, eBioscience) antibodies respectively, and removed by the magnet-mediated EasySep Cell Isolation technology (STEMCELL). The remaining cells were stained with APC-Cy7 conjugated anti-CD326 (G8.8, Biolegend) and DAPI (D9542, Sigma). Epithelial cells were sorted and collected as DAPI-;CD326(Epcam)+;tdTomato+ on a FACSAriaII high speed sorter (BD Biosciences).

### Single-cell RNA-seq and data analysis

Following FACS, single cell libraries were generated from sorted epithelial cells by using Chromium Single Cell 3’ v3 kit (10X Genomics). Sequencing was carried out on NovaSeq (Illumina) platform at IGM, UCSD. For data analysis, Cell Ranger (version 3.0.2) was used to initially align the raw reads onto the mouse reference genome (mm10) and generate the feature-barcode matrix. Next, R package Seurat (version 4.0) (Hao et al., 2021) was used to perform data quality control, normalization, scaling, principal components analysis (PCA) and the canonical correlation analysis (CCA) based data integration. Briefly, to filter out low-quality cells or doublets, cells with fewer than 200 or more than 4,000 unique features, or more than 15% mitochondrial contents were removed from further analysis. Then “LogNormalize” was used to normalize the feature expression. A total of 2,000 top variable features were identified by function FindVariableFeatures and kept for PCA. Top 20 significant components were chosen to conduct dimensional reduction by uniform manifold approximation and projection (UMAP) with default parameters. The expression features of marker genes were profiled and visualize by R package ggplot2 (version 3.3.2).

### Pathway enrichment analysis

Gene set enrichment analysis in single cell was carried out by using the R package irGSEA (https://github.com/chuiqin/irGSEA/). The enrichment of pathways in KEGG and Reactome was initially scored for individual cells by using four different gene set enrichment methods (AUCell, UCell, singscore and ssgsea). Next Wilcox test was perform to all enrichment score matrices and gene sets with adjusted p value < 0.05 are used to integrate through R package Robust Rank Aggregation (version 1.2.1) (Kolde et al., 2012). Density scatterplot of enriched pathways on the UMAP is generated by R package Nebulosa (1.0.2) (Alquicira-Hernandez and Powell, 2021).

### Influenza PR8 infection

Mouse adapted influenza strain H1N1/PR8 was obtained from ATCC (VR-95PQ) and used in all viral infection. Both male and female mice between 9-11 weeks were used. Deep mouse anesthesia was achieved by ketamine-xylazine injection. PR8 was intranasally administrated at a dose of ∼0.5 LD_50_ in a total volume of 35 μL per mouse. PBS is used for virus dilution and delivered in the same way to the control group. All procedures were conducted in the BSL2 facility at UCSD.

### Data Availability

Raw and processed files from bulk and scRNA-seq have been uploaded to the GEO database under accession number GSEXXXXXX.

## ACKNOWLEDGEMENTS

The authors would like to thank members of the Sun lab for insightful discussions. Slide scanning was performed by Olympus VS200 Slide Scanner at the UCSD School of Medicine Microscopy Core, supported by NINDS P30NS047101. Sequencing data generated at the UCSD IGM Genomics Center utilize an Illumina NovaSeq 6000 that was purchased with funding from a NIH SIG grant (#S10 OD026929). This work was supported by NIH R01HL160019 (to X.S.) and AHA postdoctoral fellowship # 827787 (to L.X.).

## AUTHOR CONTRIBUTION

L.X. and X.S. conceived and designed experiments. L.X., C.T.T., J.B. and N.T. performed experiments. L.X. analyzed bulk and single cell RNA-seq data. D.M., Y.F.S., W.K.C. and X.S. provided supervision and support. L.X. and X.S. wrote the manuscript with input from the other authors.

---

**Figure S1.**
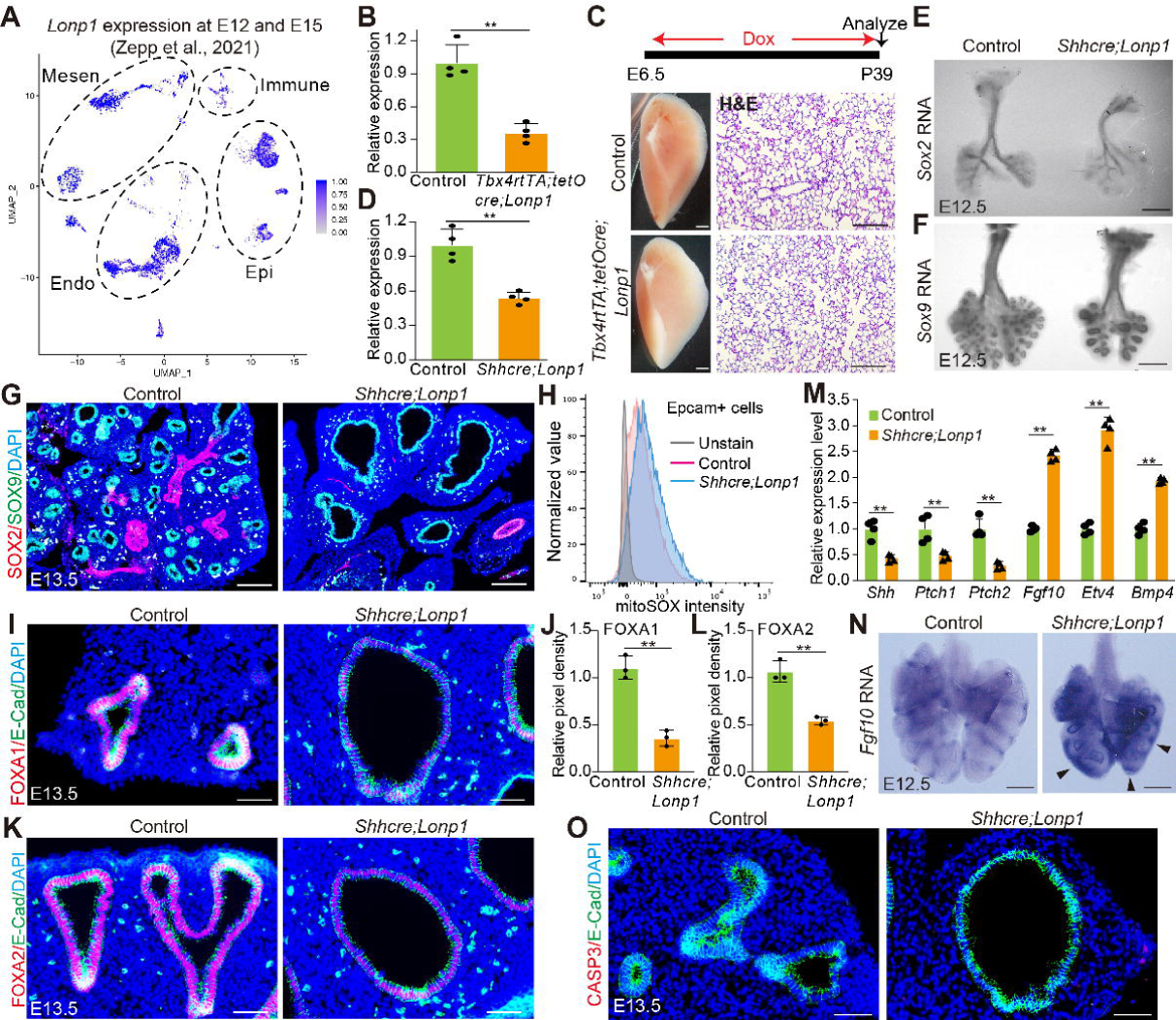
Characterization of embryonic lungs in *Lonp1* mutants and in controls. (A) *Lonp1* expression in the UMAP embedding of cells from embryonic lungs from E12.5 and E15.5 as assayed by scRNA-seq from Zepp. et al. 2021. (B) RNA levels of *Lonp1* in *Tbx4rtTA;tetOcre;Lonp1* and control at E13.5. n=4 for each group. (C) Lung morphology and H&E staining of *Tbx4rtTA;tetOcre;Lonp1* and control at P39. Scale bars, 1 mm for whole left lobe. 100 µm for H&E. (D) RNA levels of *Lonp1* in *Shhcre;Lonp1* and control at E13.5. n=4 for each group. (E) In situ hybridization for *Sox2* RNAs. Scale bars, 50 µm. (F) In situ hybridization for *Sox9* RNAs. Scale bars, 50 µm. (G)Immunofluorescent labeling of SOX2 and SOX9 proteins in *Shhcre;Lonp1* and control at E13.5. Scale bars, 100 µm. (H) Mitochondria ROS quantification by MitoSOX in *Shhcre;Lonp1* and control at E13.5, followed by flow cytometry analysis. (I-J) Immunofluorescent labeling of FOXA1 in *Shhcre;Lonp1* and control at E13.5. Quantifications were shown in (J). n=3 for each group. Scale bars, 50 µm. (K-L) Immunofluorescent labeling of FOXA2 in *Shhcre;Lonp1* and control at E13.5. Quantifications were shown in (L). n=3 for each group. Scale bars, 50 µm. (M) qPCR of representative genes in SHH and FGF10 signaling at E13.5. n=4 for each group. (N) In situ hybridization for *Fgf10* RNAs at E12.5. Scale bars, 50 µm. (O) Cleaved Caspase3 staining was not found neither in control nor in mutant naïve epithelium. Scale bars, 50 µm.

**Figure S2.**
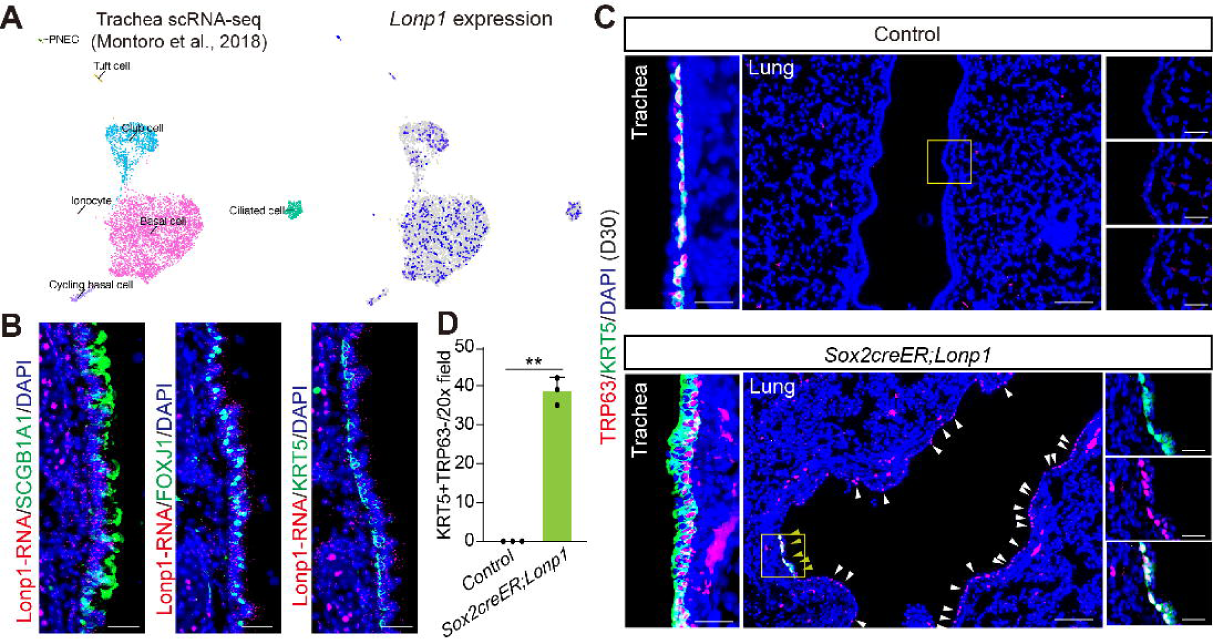
*Lonp1* is ubiquitously expressed in adult airway epithelium, loss of which led to ectopic appearance of basal-like cells in the lung. (A) *Lonp1* expression in the UMAP embedding of cells from adult trachea assayed by scRNA-seq from Montoro. et al. 2018. (B) *Lonp1* RNA detection by RNAscope, counterstained with club, ciliated and basal cell markers. Scale bars, 20 µm. (C-D) Immunofluorescent labeling of basal cells in the trachea and lung of control and *Sox2creER;Lonp1* at D30 post inactivation. The magnified regions of the ectopic basal-like cells marked by the yellow boxes in the mutant lung and the corresponding control were shown in the right. Quantifications of the luminal staining of KRT5 in trachea were shown in (D). n=3 for each group. Scale bars, 25 µm for trachea in the left, 100 µm for lung in the middle and 25 µm for zoomed pictures in the right.

**Figure S3.**
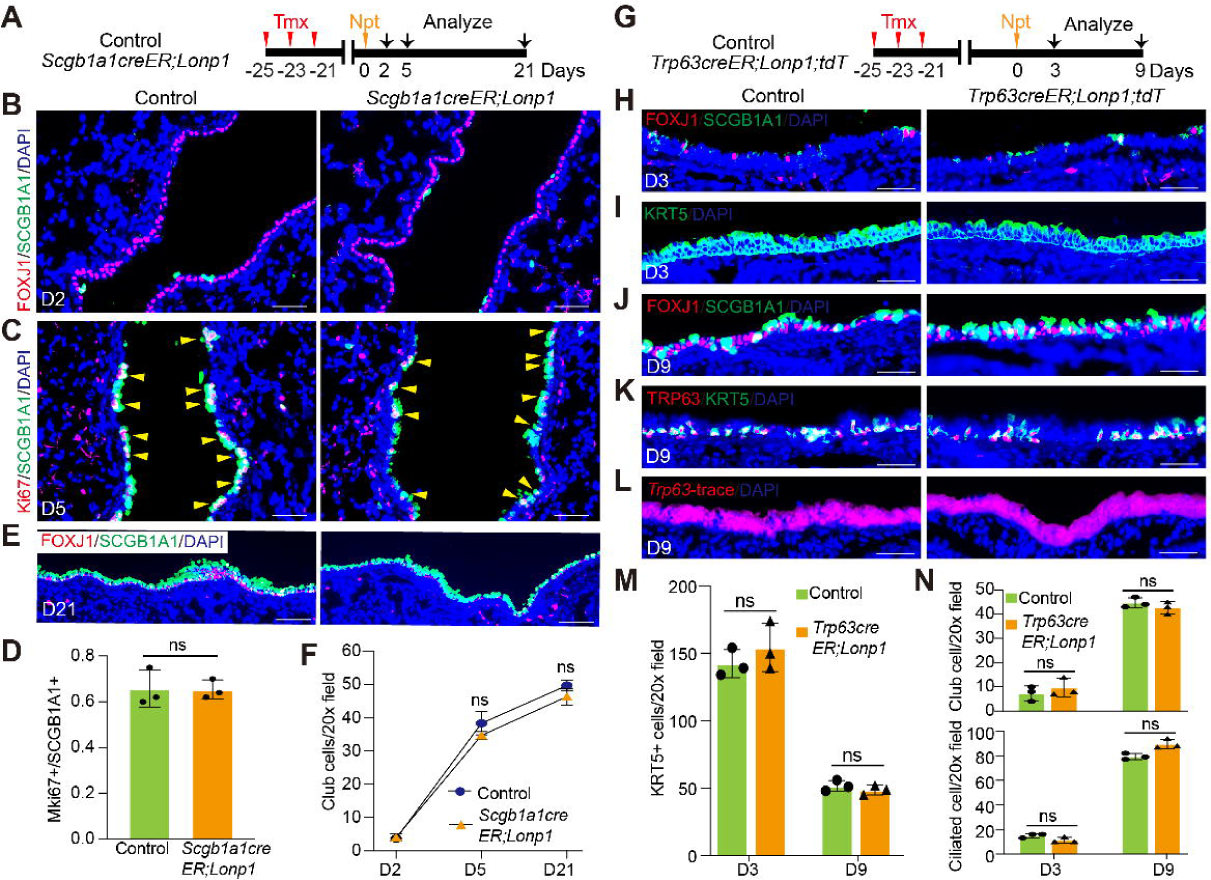
LONP1 is dispensable for the stemness maintenance of club and basal cells. (A) Schematic of experimental procedure used to inactivate *Lonp1* in club cells. (B) Most of the club cells were ablated in the lung airway 2 days post naphthalene injury. Scale bars, 50 µm. (C-D) Club cell proliferation at 5 days post injury as assayed by Mki67 staining. Quantifications were shown in (D). n=3 for each group. Scale bars, 50 µm. (E) Restoration of airway epithelium in both *Scgb1a1creER;Lonp1* and control at 21 days post injury. Scale bars, 100 µm. (F) Quantification of club cells for each time point post naphthalene injury. (G) Schematic of experimental procedure used to inactivate *Lonp1* in basal stem cells with a lineage marker. (H) Most of the club and ciliated cells were ablated in the trachea airway at 3 days post naphthalene injury. Scale bars, 20 µm. (I) Basal cell hyperplasia in *Trp63creER;Lonp1;tdT* and control trachea at 3 days post injury. Scale bars, 20 µm. (J) Regeneration of both club and ciliated cells in the trachea airway at 9 days post injury. Scale bars, 20 µm. (K) Basal cells were regressed to the basement of the trachea airway at 9 days post injury. Scale bars, 20 µm. (L) The regenerated airway epithelium was derived from basal cells as revealed by the global expression of lineage marker tdTomato. Scale bars, 20 µm. (M) Quantification of basal cells at D3 and D9 post naphthalene injury. (N) Quantification of club and ciliated cells at D3 and D9 post naphthalene injury.

**Figure S4.**
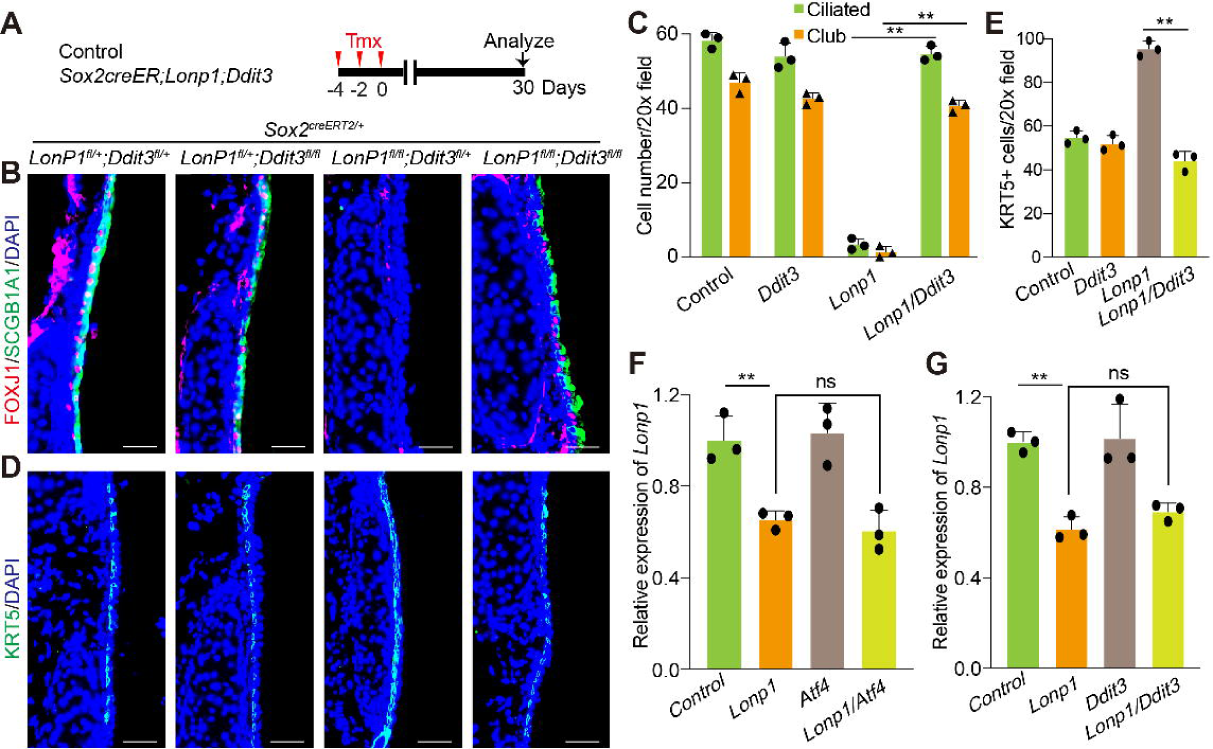
Genetic rescue of airway defects in *Sox2creER;Lonp1* by additional inactivation of *Ddit3*. (A) Schematic of experimental procedure used to analyze airway phenotypes in *Sox2creER;Lonp1;Ddit3*. (B-C) Immunofluorescent labeling of club and ciliated cells in the trachea of control and relative mutants at 30 days post inactivation. Quantifications were shown in (C). n=3 for each group. Scale bars, 20 µm. (D-E) Immunofluorescent labeling of basal cells in the trachea of control and relative mutants at 30 days post inactivation. Quantifications were shown in (E). n=3 for each group. Scale bars, 20 µm. (F) RNA levels of *Lonp1* in control, *Lonp1* or *Atf4* single mutants and *Lonp1;Atf4* double mutants as assayed by qPCR. n=3 for each group. (G)RNA levels of *Lonp1* in control, *Lonp1* or *Ddit3* single mutants and *Lonp1;Ddit3* double mutants as assayed by qPCR. n=3 for each group.

**Figure S5.**
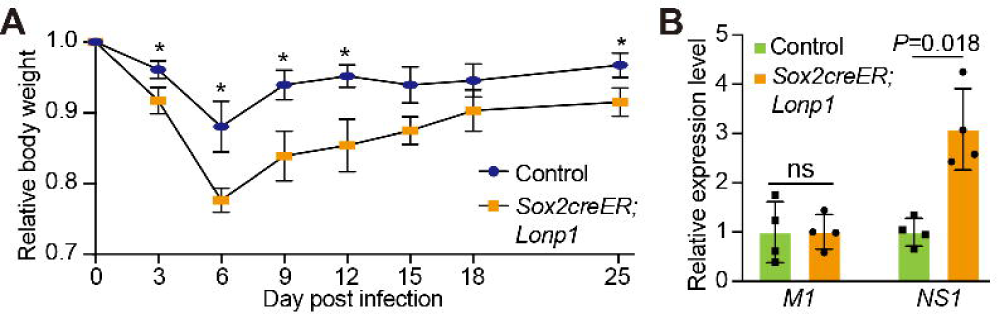
Quantification of body weight and viral loads in lungs of *Sox2creER;Lonp1* and controls post influenza infection. (A) Body weight loss of control and *Sox2creER;Lonp1* post influenza infection quantified for each time point. n=3 for each group. (B) Viral loads in control and *Sox2creER;Lonp1* lungs at 6 days post infection, as quantified by qPCR. Two pairs of primers targeting viral genes *M1* and *NS1*, respectively, were used for analysis. n=4 for each group.

